# A reversibly induced CRISPRi system targeting Photosystem II in the cyanobacterium *Synechocystis* sp. PCC 6803

**DOI:** 10.1101/2020.03.24.005744

**Authors:** Deng Liu, Virginia M. Johnson, Himadri B. Pakrasi

## Abstract

The cyanobacterium *Synechocystis* sp. PCC 6803 is used as a model organism to study photosynthesis, as it can utilize glucose as the sole carbon source to support its growth under heterotrophic conditions. CRISPR interference (CRISPRi) has been widely applied to repress the transcription of genes in a targeted manner in cyanobacteria. However, a robust and reversible induced CRISPRi system has not been explored in *Synechocystis* 6803 to knock down and recover the expression of a targeted gene. In this study, we built a tightly controlled chimeric promoter, P_*rhaBAD*-RSW_, in which a theophylline responsive riboswitch was integrated into a rhamnose-inducible promoter system. We applied this promoter to drive the expression of ddCpf1 (DNase-dead Cpf1 nuclease) in a CRISPRi system and chose the PSII reaction center gene *psbD* (D2 protein) to target for repression. *psbD* was specifically knocked down by over 95% of its native expression, leading to severely inhibited Photosystem II activity and growth of *Synechocystis* 6803 under photoautotrophic conditions. Significantly, removal of the inducers rhamnose and theophylline reversed repression by CRISPRi. Expression of PsbD recovered following release of repression, coupled with increased Photosystem II content and activity. This reversibly induced CRISPRi system in *Synechocystis* 6803 represents a new strategy for study of the biogenesis of photosynthetic complexes in cyanobacteria.

## INTRODUCTION

Cyanobacteria are oxygenic photosynthetic prokaryotes that can use light energy to fuel all intracellular biological processes. Cyanobacteria have been used as hosts to study photosynthetic processes due to their relative ease of genetic manipulation as compared to eukaryotic phototrophs. *Synechocystis* sp. PCC 6803 (hereafter *Synechocystis* 6803) has been used as a model cyanobacterial organism for decades,^1^ and many genetic tools and elements have been adapted and developed for this strain, which facilitate its genetic modification.^2^ Importantly, *Synechocystis* 6803 can use glucose as a carbon source to support heterotrophic growth, a desirable trait for the study of genes essential for photosynthesis.

Photosystem II (PSII) is a unique light-driven enzyme that oxidizes water into molecular oxygen, providing oxygen and the reducing equivalents to fix carbon, powering much of life on earth.^3–4^ PSII is an approximately 20 subunit membrane-protein complex. It binds numerous pigments and metal cofactors, as well as additional proteins that aid in assembly of the complex.^5^ PSII has a modular architecture, consisting of a reaction-center “core” module where primary charge separation occurs and two chlorophyll-binding “antenna” modules. The reaction center consists of the D1 and D2 protein subunits that bind the primary electron-transfer cofactors, along with several lower-molecular weight structural proteins. The antenna modules are based around the chlorophyll-binding protein subunits CP43 and CP47 and associated lower molecular weight structural proteins. A catalytic tetra-manganese cofactor is liganded primarily by the D1 protein, with one ligand supplied by the CP43 protein.

PSII, due to its demanding photochemistry, undergoes frequent photo-oxidative damage and must be assembled and repaired in an efficient and tightly regulated manner to maintain high photosynthetic activity.^6–12^ The D1 and D2 reaction center proteins are the most frequently damaged, as they are closest in proximity to the high-energy electron-transfer reactions taking place.^12–14^ To repair the damaged complex while maintaining the undamaged and expensive-to-synthesize chlorophyll-protein antenna modules, D1 and D2 are selectively removed and replaced by newly synthesized protein copies. This process of continual assembly and repair is called the PSII lifecycle, and has been extensively studied in mutants lacking key proteins in photosynthetic processes that are halted at various stages in PSII assembly.^15–18^ However, the lifecycle has been difficult to study as a complete cycle in the same organism due to the low abundance of intermediate complexes in photosynthetic cells, and the simultaneous presence of various PSII intermediate complexes at different stages of the lifecycle CRISPR/Cas (clustered regularly interspersed palindromic repeats/CRISPR-associated protein) technology has revolutionized the biotechnology field as a technique to precisely edit both eukaryotic and prokaryotic genomes, including cyanobacteria.^19–22^ As a complement to CRISPR/Cas technology, CRISPR interference (CRISPRi) utilizes a nuclease-deficient Cas protein (dCas protein) to bind to a target sequence but not introduce a double-stranded break. A CRISPRi system contains a dCas protein and a guide RNA (gRNA), which together form a dCas-gRNA ribonucleoprotein complex that binds to a genomic site complementary to the gRNA sequence. By binding to its target site, the dCas-gRNA complex blocks transcription and inhibits the expression of specific genes.^23^ CRISPRi, utilizing either dCas9 or ddCpf1 (DNase-dead Cpf1, also known as dCas12a), has been applied to several species of cyanobacteria.^19, 24–28^ However, only dCas9-CRISPRi systems have been tested for regulation of genes in *Synechocystis* 6803.^24, 29^

To dynamically and precisely regulate the expression of target genes, a CRISPRi system driven by a tightly controlled inducible promoter is of great interest. For *Synechocystis* 6803, many inducible promoters have been characterized,^30–32^ but only the anhydrotetracycline (aTc)-inducible promoter has been applied to CRISPRi in *Synechocystis* 6803.^33^ This TetR-regulated inducible system is challenging because aTc is light degradable, which leads to unreliable performance under phototrophic conditions. The rhamnose responsive promoter *P_rhaBAD_* has been shown to be a robust and inducible system in *Synechocystis* 6803. This promoter uses the *E. Coli* transcription factor RhaS as the regulator.^32, 34^ In this study, we used and modified the *P_rhaBAD_* promoter to drive expression of the ddCpf1 protein for CRISPRi, as the ddCpf1 protein has been shown to have lower toxicity in cyanobacteria than dCas9.^21–22^ We added a riboswitch to construct a tightly controlled promoter cassette that showed titratable behavior over a broad range of expression. When this inducible promoter is applied to our CRISPRi system, expression of the targeted gene *psbD* (encoding for D2 protein of Photosystem II) is reduced by over 95% with inducers added, while no repression was observed without inducers. Importantly, recovery of D2 expression was detected after release of CRISPRi repression by washing out inducers. We knocked down key PSII proteins through CRISPRi to almost eliminate formation of Photosystem II and then recovered the formation of the photosystem in its native milieu. Our system is an invaluable resource to study the PSII life cycle in a targeted manner and represents a new framework for studying *de novo* assembly of photosystems.

## RESULTS AND DISCUSSION

### Repression of *psbD* genes in *Synechocystis* 6803 by CRISPRi

The plasmid pCRISPR i-D2 was constructed, using *P_rhaBAD_* to drive transcription of both *ddcpf1* and a gRNA targeted to the *psbD* genes (Figure 1A and Table S1). PsbD (D2) is a structural part of the core PSII reaction center,^18^ and essential for photoautotrophy. In *Synechocystis* 6803, there are two *psbD* genes, *psbD1* and *psbD2*, which encode identical proteins but have slightly different nucleotide sequences. The site targeted on both *psbD* genes has the same sequence in both (the PAM sequence TTG is 7 nucleotides behind the start codon), so both *psbD* genes will be targeted by the same gRNA. Furthermore, *psbD* shares an operon with and is directly upstream of the *psbC* gene, which codes for the CP43 protein (Figure 1A), so with one gRNA, we expect to observe repression of both *psbD* genes and *psbC*.

**Figure 1.**
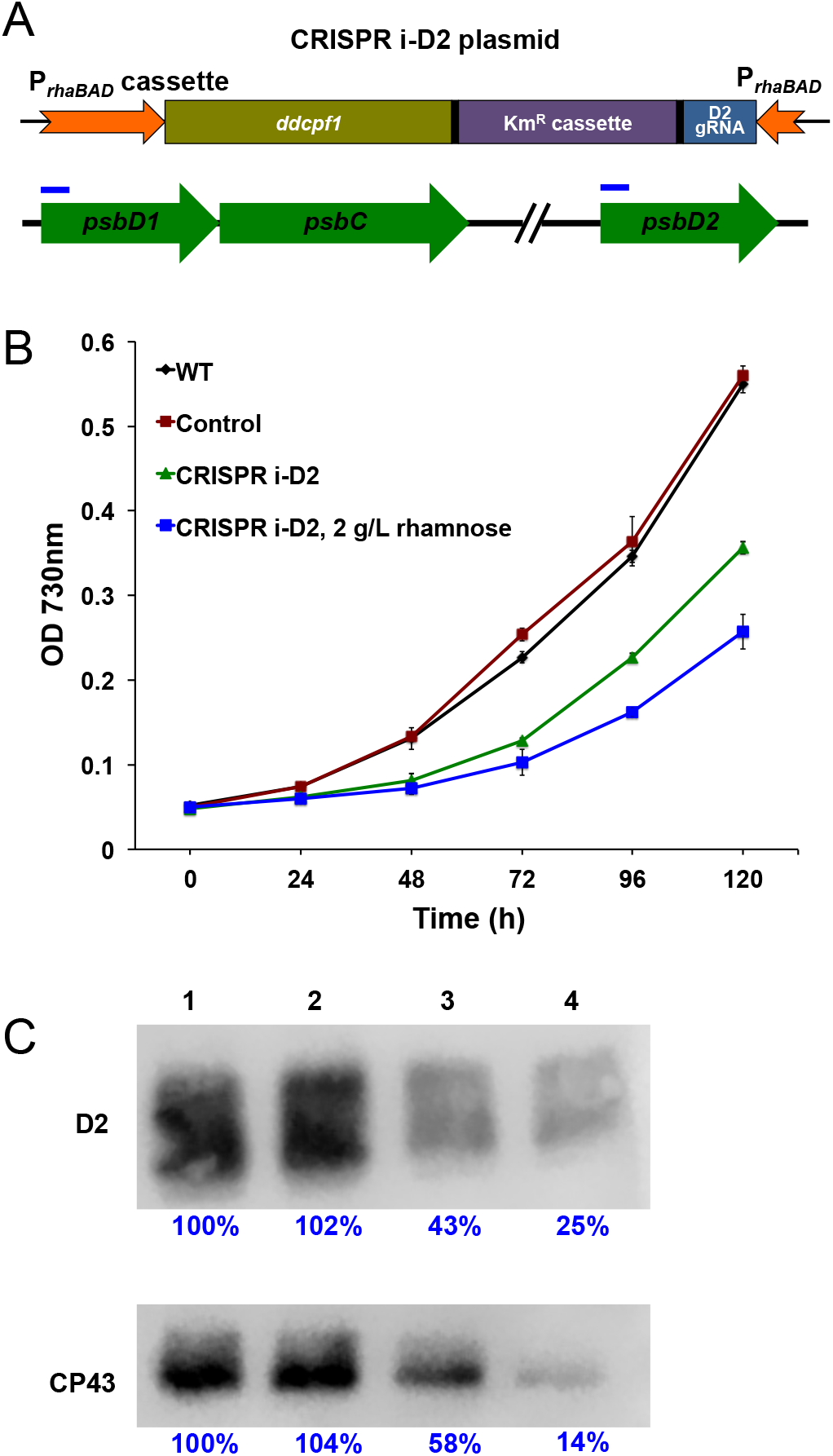
Repressing the expression of *psbD* gene in *Synechocystis* 6803 through CRISPRi. (A) Diagram of CRISPRi system on the replicating plasmid pRSF1010. Promoters are shown as orange arrows, while *ddcpf1* gene and gRNA part are yellow and blue rectangles, respectively. The targeting sites of gRNA to the psbD genes are indicated by blue bars. (B) Growth curves of *Synechocystis* 6803 strains cultured in BG11 medium, shaking in flask under 30°C with light intensity 30 μmol photons m^-2^ s^-1^. CRISPR i-D2 is the strain containing the CRISPRi plasmid with gRNA targeting to *psbD* genes. Control is the strain containing the CRISPRi plasmid but with the gRNA targeting to the *eyfp* gene. 2 g/L rhamnose was added to the culture as the inducer to promote the transcription of *ddcpf1* and gRNA. Error bars represent the standard deviations observed from at least three independent experiments. (C) Expression levels of D2 and CP43 detected by western blot. Lane 1 to 4 represent loading 20 μg of total membrane proteins extracted from WT, control, CRISPR i-D2, and CRISPR i-D2 with 2 g/L rhamnose, respectively. Blue percentages are the relative blot intensities for each protein band.

After transfer of pCRISPR i-D2 (Figure S1) into wild type (WT) of *Synechocystis* 6803, we observed growth under autotrophic conditions (Figure 1B). WT and the CK01 control strain (CRISPRi plasmid with gRNA targeted to *eyfp)* grew similarly. However, growth of CRISPRi-D2 was impaired compared to WT, indicating that PSII is impaired in this strain even under un-induced conditions, and that *P_rhaBAD_* drives expression of ddCpf1 even without the addition of rhamnose. Conversely, in the cells induced with 2 g/L rhamnose, growth was not completely abolished, indicating that there is not full repression of PSII genes by CRISPRi in the induced strain, even though the growth rate is slower than the culture without rhamnose (Figure 1B). Intracellular protein levels of D2 and CP43 as measured by western blot were consistent with growth phenotypes. About 50% of WT D2 and CP43 protein levels were detected in the uninduced CRISPR i-D2 strain (Figure 1C) and about 20% of D2 and CP43 protein levels were observed in the CRISPR i-D2 strain with added rhamnose. These results show that the P_*rhaBAD*_ promoter is not tightly regulated nor is it strong enough to support full repression.

To confirm that the *P_rhaBAD_* promoter has leaky expression, we placed the *P_rhaBAD_* cassette in front of the EYFP coding gene in *Synechocystis* 6803 to make the plasmid pRhaBAD-eyfp (Figure S2). EYFP fluorescence was compared to a control strain CK3068,^35^ in which the promoter in front of *eyfp* is removed. We observed a 2 to 2.5-fold higher level of fluorescence over the CK3068 strain in the uninduced culture (Figure S2). This directly confirmed that the *P_rhaBAD_* cassette as utilized here does not have tight control over expression of downstream genes. A reason might be the promoter *P_lacOI_* we use to drive the regulator protein RhaS in the cassette (Figure S1). RhaS is the regulator for the rhamnose responsive system, and its expression level is a key factor for the RhaBAD promoter to control downstream gene(s).^36^ The promoter used here might lead to an unbalanced intracellular level of the RhaS protein as compared with that used by Kelly et. al.^34^

Ideally, the CRISPRi system would maintain full photosynthetic activity under uninduced conditions, while being repressed nearly 100% in an induced condition, to become photoautotrophic and non-photoautotrophic in each state, respectively. This would allow us to observe complete *de novo* assembly of Photosystem II.

### Construction of a chimeric promoter optimized the performance of the *P_rhaBAD_* promoter

To tighten the control and improve the strength of the promoter *P_rhaBAD_*, we switched out its ribosome binding sequence (RBS) for an RBS containing the theophylline responsive riboswitch (RSW), which has been effective in controlling the translation of downstream genes in cyanobacteria.^37–38^ We used the riboswitch sequence ‘E’ from Ma et. al.^37^ (Table S1). We call this new promoter *P*_*rhaBAD*-RSW_ and placed the EYFP coding gene under its control to make the P_*rhaBAD*-RSW_-EYFP strain (Figure 2A).

**Figure 2.**
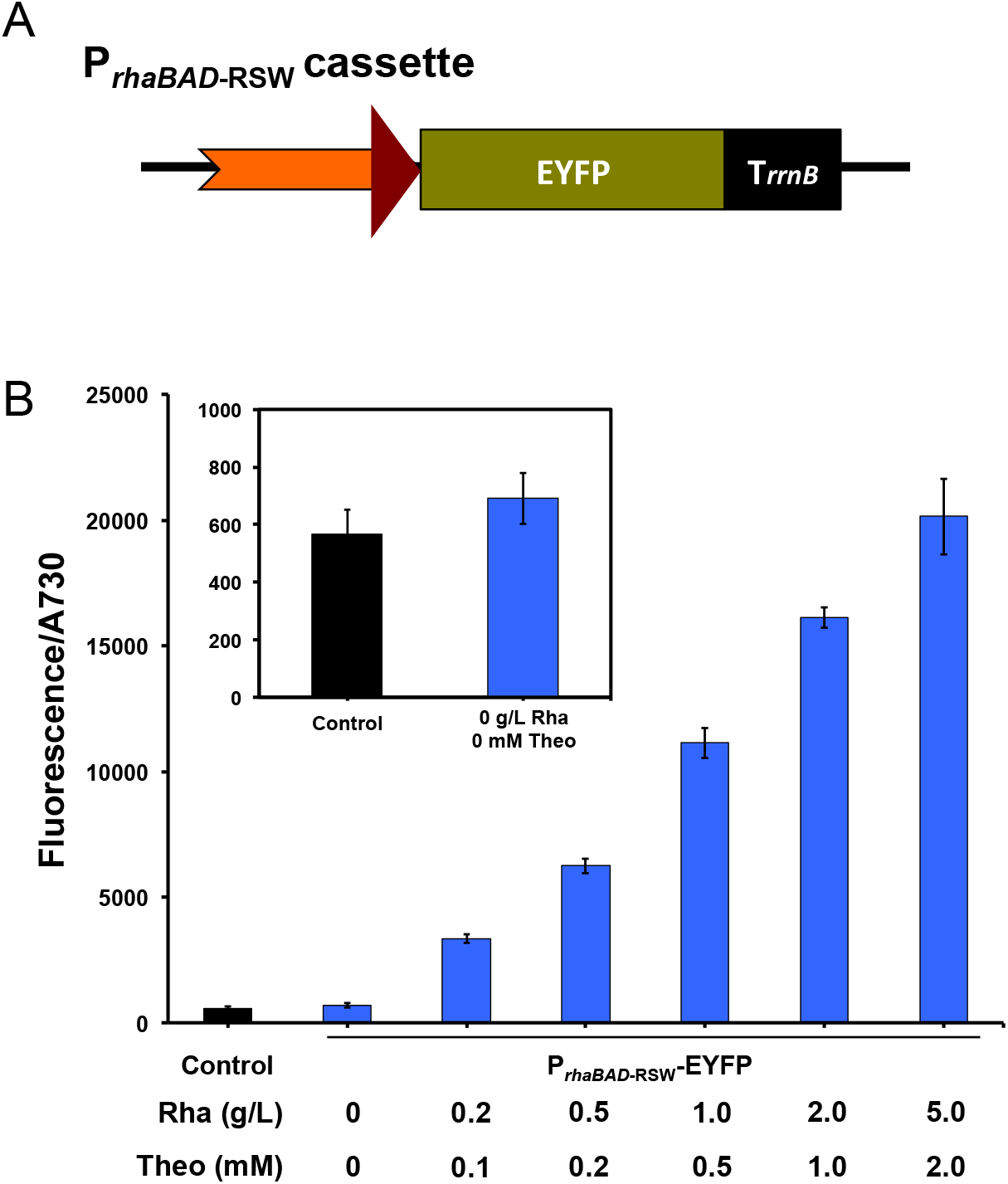
Adding theophylline responsive riboswitch (RSW) optimizes the inducible promoter system. (A) Diagram of the P_*rhaBAD*-RSW_ cassette driving the expression of EYFP on the replicating plasmid pRSF1010. The riboswitch is shown in red triangle, switching the RBS of promoter P_*rhaBAD*_, which is shown in orange arrow. (B) Expression of reporter gene coding for EYFP driven by *P*_*rhaBAD*-RSW_. Fluorescence data were normalized to optical density at 730 nm. Control is the strain containing the plasmid with EYFP encoding gene but without any promoters to drive it. Rhamnose (Rha) and theophylline (Theo) were added to the culture with the concentrations as indicated. The inset panel magnifies the control and un-induced strains. Strains were cultured in BG11 medium, shaking in flask under 30°C with light intensity 30 μmol photons m^-2^ s^-1^. Samples were collected after 96 hours culture for EYFP assay. Error bars represent the standard deviations observed from at least three independent experiments.

Cell samples of P_*rhaBAD*-RSW_-EYFP strain were collected for fluorescence assay after 96 hours of growth in BG11 medium under various inducer concentrations. Fluorescence of the culture without inducers was not significantly different from that of the control strain CK3068 (Figure 2B), which indicated that there is not leaky expression from P_*rhaBAD*-RSW_. Fluorescence intensity increased with the concentration of inducers up to 5 g/L rhamnose and 2 mM theophylline, with a 120-fold difference between the uninduced culture and the culture with the highest concentrations of inducers (Figure 2B). Additionally, we tested the effect of adding either inducer at a time on the expression of the *eyfp* gene. We observed that adding only rhamnose achieved 90% of the fluorescence level as adding both inducers (Figure S3), indicating that activation by the chimeric promoter occurs mainly on the transcriptional level, and the riboswitch primarily prevents un-induced expression. All above results suggested that P_*rhaBAD*-RSW_ is a tightly controllable and titratable promoter cassette with a wide induction range. We applied P_*rhaBAD*-RSW_ to drive the expression of *ddcpf1* in our CRISPRi system in subsequent experiments.

### Tightly controlled expression of D2 and CP43 by the CRISPRi_RSW_ system

We constructed a new plasmid applying the P_*rhaBAD*-RSW_ promoter to control the expression of *ddcpf1* and containing the same gRNA sequence targeting to *psbD* genes, named as the pCRISPR i_RSW_-D2 plasmid (Figure 3A). After transfer into WT *Synechocystis* 6803, the CRISPR i_RSW_-D2 strain grew similarly to WT under photoautotrophic conditions without inducers. When 5 g/L rhamnose and 2 mM theophylline were added, growth of the CRISPR i_RSW_-D2 strain was severely inhibited under autotrophic conditions (Figure 3B). This growth phenotype indicates that PSII is fully active in the un-induced strain, while has low to no activity when the strain is induced. To further analyze CRISPR i_RSW_-D2, subsequent experiments are performed with the addition of 10g/L glucose to the growth medium.

**Figure 3.**
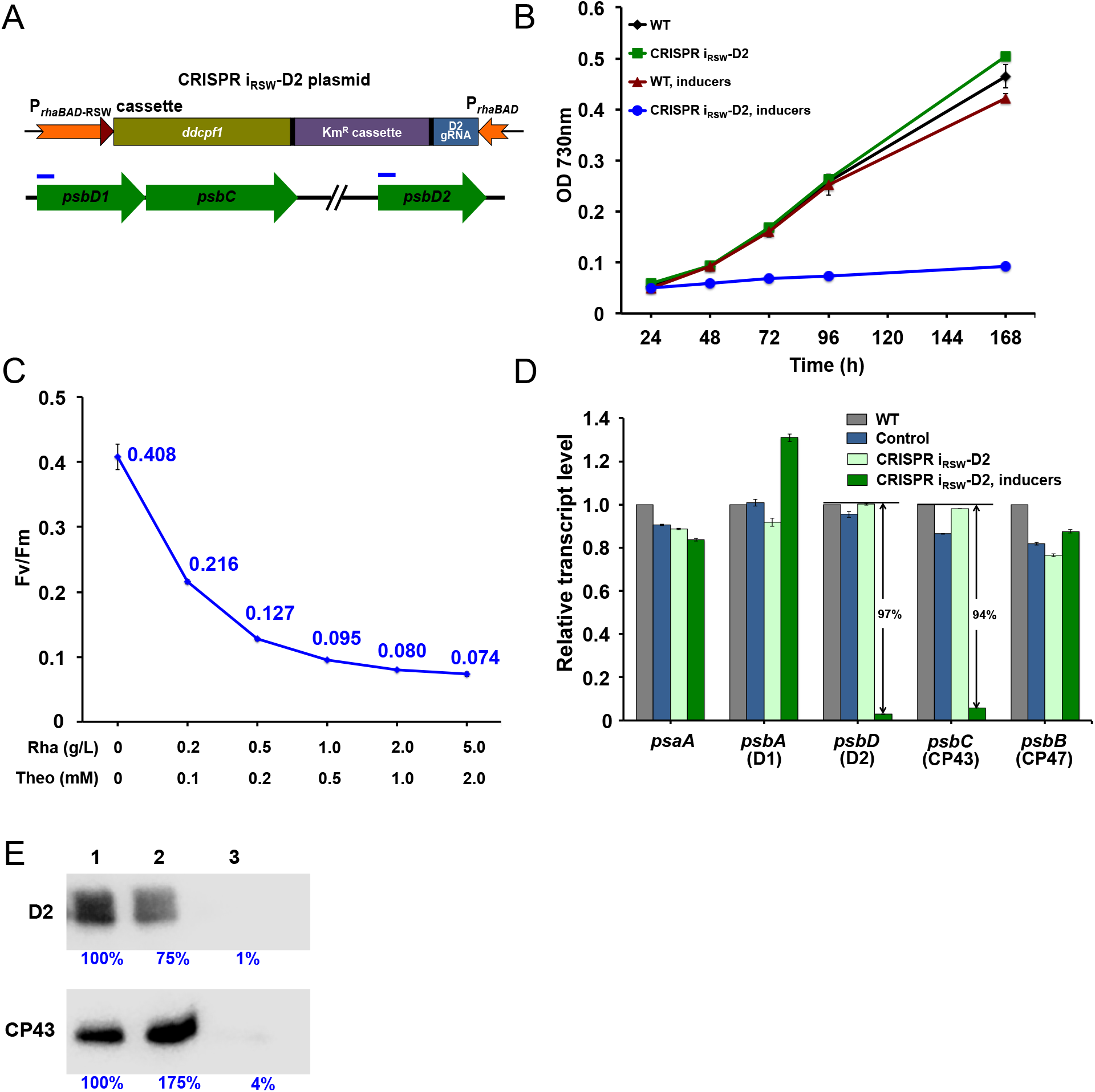
Repressing the expression of D2 and CP43 encoding genes by the inducible CRISPR i_RSW_ system. (A) Diagram of CRISPR i_RSW_ system on the replicating plasmid pRSF1010. Promoters are shown in orange, the riboswitch in red triangle, and *ddcpf1* gene and gRNA are in yellow and blue, respectively. The targeting sites of gRNA to the *psbD* genes are indicated by blue bars. (B) Growth curves of *Synechocystis* 6803 strains cultured in BG11 medium. The strain containing the CRISPR i_RSW_ system with the gRNA targeting to *psbD* genes was labeled as CRISPR i_RSW_-D2. (C) The Fv/Fm values correspond to inducer concentrations. Rhamnose (Rha) and theophylline (Theo) were added to the culture with the concentrations indicated. All strains were cultured in BG11 medium with 10 g/L glucose and sampled for Fv/Fm assay when OD 730nm reached 1.0-1.5. (D) Transcriptional levels of photosystem genes quantified by q-PCR. Cell samples were collected from the culture in BG11 medium with 10 g/L glucose when OD 730nm reached 1.0-1.5, and total RNA was extracted for analysis. (E) Protein levels of D2 and CP43 detected by western blot. Lane 1 to 3 represent loading 20 μg of total membrane proteins extracted from cell samples of control, CRISPR i_RSW_-D2, and CRISPR i_RSW_-D2 with inducers, respectively. Blue numbers are the relative blot intensities of corresponding proteins. Cell samples were collected when OD 730nm reached 1.0-1.5 from the culture in BG11 medium with 10 g/L glucose. Error bars in the figure represent the standard deviations observed from at least three independent experiments. Except where notified, 5 g/L rhamnose and 2 mM theophylline were added as inducers to the culture. Control is the strain containing the CRISPR i_RSW_ plasmid but with the gRNA targeting to the *eyfp* gene. Cells are cultured in flask shaking under 30°C with light intensity 30 μmol photons m^-2^ s^-1^.

If PSII content is reduced by the CRISPRi_RSW_ system, then we expect PSII activity to be lower in cells cultured with inducers than without inducers. We used a PAM fluorometer to assay the F_V_/F_M_, or PSII quantum yield, of cultures. The ratio F_V_/ F_M_ is a relative measure of PSII activity in cells, and the higher the value, the more functional PSII is present. F_V_ (variable fluorescence) represents the difference between F_m_ (maximal fluorescence) and F_0_ (background fluorescence) and can be attributed to fluorescence arising directly from active PSII reaction centers.^39^ The F_V_/F_M_ of the CRISPR i_RSW_-D2 strain decreased with increased concentrations of inducers (Figure 3C). At the highest concentration of inducers, 5 g/L rhamnose and 2 mM theophylline, the FV/FM is 18% of that of the un-induced culture. The above results indicate that, consistent with control of EYFP by CRISPRi_RSW_, control of PSII content is titratable by our CRISPRi_RSW_ system.

To check the specificity of CRISPR i_RSW_-D2 and quantify the level of repression, both semi-quantitative RT-PCR and quantitative PCR (q-PCR) were performed on WT, the CK02 control strain (containing the CRISPRi_RSW_ plasmid with gRNA targeted to *eyfp*), and the CRISPR i_RSW_-D2 strain. In addition to the targeted genes *psbD* (D2) and *psbC* (CP43), the Photosystem I structural gene *psaA* and PSII reaction center genes *psbA* (D1) and *psbB* (CP47) were assayed as transcriptional controls. Both RT-PCR and q-PCR showed that no difference was observed between the three strains under photoautotrophic, uninduced conditions for all 5 genes (Figure S4 and S5). Because the CRISPR i_RSW_-D2 strain does not grow when induced under photoautotrophic conditions in BG11, samples were collected only from WT, CK02, and the un-induced CRISPR i_RSW_-D2 strain after 7 days growth under this condition.

However, when the strains were grown in BG11 medium with glucose, the *psbD* and *psbC* genes in the induced CRISPR i_RSW_-D2 strain showed 97% and 94% decrease in transcription, respectively (Figure 3D and Figure S4). Transcription of the *psaA* gene and the *psbB* gene (CP47) of PSII were unchanged, indicating a specific transcriptional response to induction of CRISPRi. Interestingly, *psbA*, which codes for the PSII protein D1, was up-regulated around 30% under induced conditions, indicating that its transcription is up-regulated in response to severely inhibited photosynthesis as a compensation mechanism, as the D1 protein is the most frequently damaged and replaced under normal and photo-inhibitory conditions.^12^ Intracellular protein levels of D2 and CP43 as measured by western blot are consistent with the mRNA levels. D2 and CP43 are reduced to 1% and 4% of un-induced levels, respectively, in the induced samples (Figure 3E).

The above results show that our CRISPRi_RSW_ system is a powerful tool to tightly and specifically control the expression of target genes over a broad range and in a titratable way in *Synechocystis* 6803. Because there is no expression of ddCpf1 without inducers, and expression of target genes is reduced by over 90% with inducers added, it is an ideal system to study the recovery process after release of repression.

### Release of CRISPRi repression allows recovery of the expression of D2 and CP43 proteins

The CRISPRi_RSW_ system effectively knocks down D2 and PSII activity and is tightly regulated. However, to study Photosystem II assembly we require relief of this inhibition, to allow PSII to rebuild following repression. We did further experiments to observe the recovery of targeted genes after release of the CRISPRi induction. To test whether PSII can fully recover in the CRISPR i_RSW_-D2 strain, the CRISPR i_RSW_-D2 strain was cultured in BG11 medium with glucose and inducers. When the OD730 nm reached 2.0, cells were washed twice and re-suspended in fresh BG11 medium at a 20-fold dilution to OD730 nm of 0.1 (time point 0). During the recovery process, cells were collected to verify the *de novo* synthesis of D2 and CP43 proteins, as well as PSII.

After release of repression, we first checked the transcription of *psbD* and *psbC* through q-PCR. Samples were collected over a time course, and we observed the gradual increase of transcript levels with time, reaching around 40% of the WT mRNA level after 8 hours and 100% after 96 hours (Figure 4A). Two phases of recovery were observed. In the first phase, the initial 24 hours, transcript level increases from 5% to 50% of WT level. This is followed by a second, slower phase over the next 72 when transcript levels increase back to 100%. We also monitored the F_V_/F_M_ increase over time (Figure 4B) following release of inhibition, which showed accumulation of active PSII in cells. Two phases of recovery were also observed for F_V_/F_M_. The first phase occurs over day 0 to day 3, during which the F_v_/F_m_ value increased from 0.07 to 0.18, while in the second phase F_V_/F_M_ increased slowly and linearly up to 0.35 over the following 13 days (Figure 4B).

**Figure 4.**
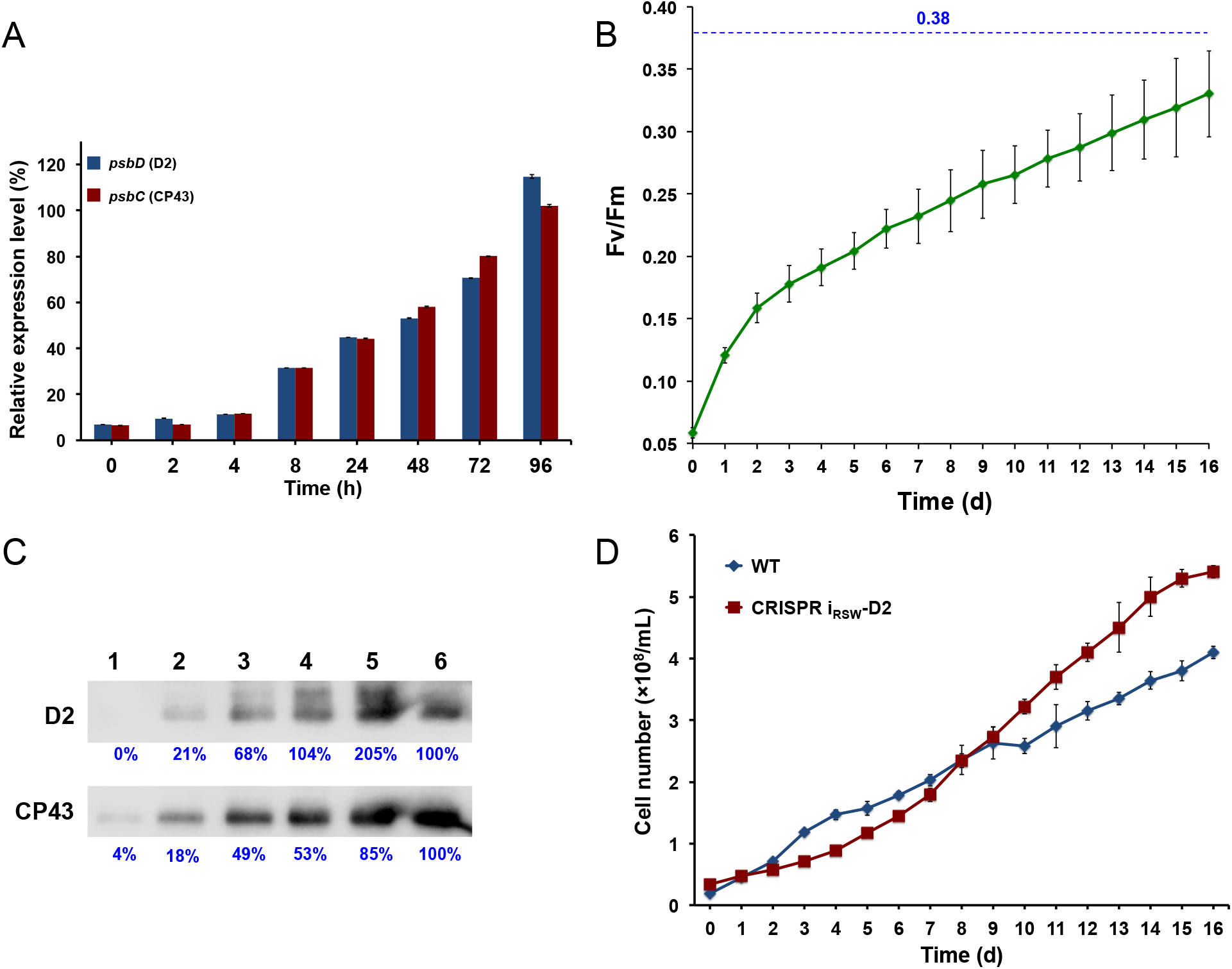
Recovery the expression of D2 and CP43 after release the repression by the CRISPR i_RSW_ system. Both WT and CRISPR i_RSW_-D2 strains were initially cultured in BG11 with glucose and inducers. After 4 days growth, cells were collected, washed twice, and re-suspended in fresh BG11 medium for monitoring. (A) Recovery of *psbD* and *psbC* transcript levels in the CRISPR i_RSW_-D2 strain determined by q-PCR. Transcript levels of both genes in WT were set as 100%. (B) The Fv/Fm values of the CRISPR i_RSW_-D2 strain increased along with time. The dotted blue line is the Fv/Fm value of WT after 16 days culture in BG11. (C) Recovery of D2 and CP43 protein levels in the CRISPR i_RSW_-D2 strain determined by western blot. Lane 1 to 5 represent 20 μg of total membrane proteins extracted from CRISPR i_RSW_-D2 when the F_v_/F_m_ value is 0.07, 0.16, 0.25, 0.35, and 0.39, respectively. Lane 6 represents 20 μg of total membrane proteins extracted from WT. Blue numbers are the corresponding blot intensities of each band. (D) Cell numbers of both strains along with time during the recovery process. Error bars in the figure represent the standard deviations observed from at least three independent experiments. Cells are cultured in flask shaking under 30°C with light intensity 30 μmol photons m^-2^ s^-1^.

Western blots (Figure 4C) showing protein recovery with increasing F_V_/F_M_ levels demonstrate a slow increase in CP43 and D2 protein levels over time. At time point 0, corresponding to F_V_/F_M_ of 0.07, D2 and C43 are less than 5% of the WT level. Interestingly, D2 protein levels increase to 200% of wild type as recovering cells approach the PSII activity of WT, while CP43 increases to just 85% of its wild type protein level. This creates a situation in which there are “extra” D2 copies. These copies of D2 could be in PSII assembly intermediate complexes, as *de novo* PSII assembly is thought to occur step-wise, first incorporating the D1 and D2 proteins, and then later the CP43 and CP47 proteins.^40^

Under photoautotrophic conditions, photosynthesis is the sole energy source for intracellular biological processes in *Synechocystis* 6803, including cell division. We expect cell division rate to be affected for the CRISPR i_RSW_-D2 strain over the recovery process. Cell numbers were counted every 24 hours during recovery for both WT and the CRISPR i_RSW_-D2 strain. As shown in Figure 4D, the two strains showed divergent cell division rate phenotypes. Compared to the WT strain, which showed more rapid division over the first 4 days before slowing, the CRISPR i_RSW_-D2 strain showed a lag phase before day 4, at which time point cells began to divide at a faster rate. This 4-day lag before division begins suggests that cell division was inhibited because of the insufficient energy generated from the low level of intracellular PSII, and correlates to the phases of the F_V_/F_M_ curve (Figure 4B). The initial rapid increase in F_V_/F_M_ can be explained by the fact that the F_V_ (or the difference in fluorescence due to PSII activity compared to background chlorophyll fluorescence) increases due to the small new amount of active PSII, while the total background chlorophyll fluorescence (F_0_) does not start to increase until the cells begin to divide. Once cells begin to grow and divide, photosystem I-associated chlorophyll increases, as does synthesis of unassembled PSII subunits containing chlorophyll, which contribute to F0, making the subsequent increase in F_V_/F_M_ slower.

D2 and CP43 are essential proteins in the core reaction center of PSII, and 18 inhibiting their biosynthesis eliminates the formation of PSII.^18^ As shown in Figure 4, a two-phase recovery was observed for all parameters measured, including transcription of *psbD* and *psbC* genes, FV/FM values, and cell numbers. This two-phase phenomenon is correlated with the PSII content, which affects the energy supply for biological processes, such as transcription, translation, and cell division. Because the CRISPRi_RSW_ system constructed here knocks down the expression of targeted genes by over 90%, the intracellular PSII activity is very low, around 15% of WT based on F_V_/F_M_ (Figure 3C and 4B), and less than 5% based on D2 and CP43 protein levels (Figure 3E and 4C). The CRISPR i_RSW_-D2 strain grew under near photoheterotrophic conditions when glucose and inducers were added to the medium. After resuspension in BG11 medium, cells must convert to autotrophic growth but with a very low initial PSII content. So, there is a rapid initial increase in mRNA and F_V_/F_M_, but energy stores are quickly used up, inhibiting cell division until sufficient energy derived from the small amount of initial PSII has allowed cells to divide and synthesize more PSII to make more energy. Therefore, there is a subsequent slow recovery of PSII activity, mRNA, and protein levels.

To study the process of PSII biogenesis, deletion of target genes is the classical genetic strategy. Using a deletion strategy, intermediate PSII assembly complexes form in the specific gene knockout mutant, and these complexes are analyzed to infer the PSII biogenesis process.^41^ The function of numerous proteins in PSII biogenesis has been identified through this strategy. However, deletion mutants cannot recover the expression of the deleted gene, so the biogenesis process cannot be systematically studied. The CRISPRi technology supplies an alternative and valuable way to uncover the biogenesis process. With this tightly controlled CRISPRi_RSW_ system in *Synechocystis* 6803, we have developed a new platform to study not only PSII biogenesis, but also assembly of other protein complexes such as Photosystem I.

## CONCLUSIONS

We built a chimeric promoter system, P_*rhaBAD*-RSW_, which exhibits tight control and a broad, titratable inducible range in *Synechocystis* 6803. Expression of ddCpf1 driven by this chimeric promoter in a CRISPRi system effectively and reversibly represses the expression of the targeted genes, *psbD* and *psbC*, and nearly inhibits the formation of PSII protein complex. Following release of the repression by removing inducers, PSII is re-built in the same cells. In contrast to a strategy of knocking-out to study gene function, our CRISPRi_RSW_ system does not modify or edit the genome. Furthermore, we supply a new platform to study the biogenesis processes, during which protein complex intermediates can be tracked dynamically following release of inhibition.

## METHODS AND MATERIALS

### Strains and culture conditions

All cloning work was performed in *E. coli* strain XL1-Blue grown in LB medium in culture tubes or on agar plates at 37 °C, supplemented with 50 μg/ml kanamycin, 20 μg/ml chloramphenicol, or 100 μg/ml ampicillin, as needed. *Synechocystis* 6803 cells were cultured in BG11 medium supplemented with 30 μg/ml kanamycin as needed, under continuous white light at 30 μmol photons m^-2^ s^-1^ at 30 °C. Cultures were grown in 125-ml glass flasks or on agar plates. To monitor growth, 150 μL of cells suspension was loaded onto a 96-well plate and OD at 730 nm was measured on a plate reader (BioTek, VT).

### Construction of recombinant plasmids and engineered strains

Plasmids and strains used in this study are listed in Table S2, and all primers used in this study are listed in Table S3. All plasmids were constructed by Gibson Assembly strategy,^42^ using linear fragments purified from PCR products. All plasmids used in this study are based on the broad host replicating plasmid pRSF1010.^43^ The DNA fragments used to assemble the CRISPRi plasmid were amplified from other plasmids in the Pakrasi lab.

A tri-parental conjugation method was used to transfer all pRSF1010 derivative plasmids to *Synechocystis* 6803 wild-type cells,^44^ using a helper strain of *E. coli* containing the pRL443 and pRL623 plasmids.^45–46^ Transformants were isolated on BG11 agar plates containing 20 μg/mL kanamycin, as needed. Isolated *Synechocystis* 6803 transformants were checked by PCR to confirm presence of the desired constructs.

All PCR amplifications were performed using Phusion High-fidelity DNA polymerase (Thermo Scientific). Plasmids and PCR products were purified using the GeneJET (Thermo Scientific) plasmid miniprep kit and gel extraction kit, respectively. Oligonucleotides were designed using the SnapGene software (GSL Biotech LLC) and synthesized by IDT (Coralville, IA). The sequences of all the plasmids constructed in this study were verified (Genewiz, NJ).

### Measurement of EYFP fluorescence

EYFP fluorescence was measured directly from cells suspension in culture. The fluorescence intensity and the optical density of each culture were determined in 96-well black-walled clear-bottom plates on a BioTek Synergy Mx plate reader (BioTek, Winooski, VT). The excitation and emission wavelengths were set to 485 nm and 528 nm for EYFP. All measured fluorescence data were normalized by culture density.

### Reverse transcription-PCR (RT-PCR) and quantitative PCR (q-PCR)

Total RNA samples for RT-PCR and q-PCR were extracted using the reagent RNAwiz (Ambion). After quantification of RNA, 200 ng of DNase-treated RNA samples, Superscript II reverse transcriptase, and random primers (Invitrogen) were used for reverse transcription according to the manufacturer’s instructions. cDNA generated after reverse transcription was used as the template for PCR to validate the transcription of genes.

QRT-PCR Sybr green dUTP mix (ABgene) was used for the q-PCR assay on an ABI 7500 system (Applied Biosystems). Each reaction was performed in triplicate, and the average threshold cycle (CT) value was used to calculate the relative transcriptional levels for the amounts of RNA. All primers used for RT-PCR and q-PCR are listed in Table S3.

### Western blot analysis

Cyanobacterial cells were harvested and broken by bead beating as described previously^47^ with minor modifications. Cells were re-suspended in RB buffer (25% glycerol (wt/vol), 10mM MgCl_2_, 5mM CaCl_2_, 50 mM MES buffer pH 6.0) and broken by vortexing with 0.17 mm glass beads. Membrane fraction was isolated by centrifugation, re-suspended in RB, and solubilized by addition of β-D-dodecyl maltoside (DDM) to a final concentration of 0.8%. After incubation on ice in dark for 30 min, the solubilized membranes were separated from the insoluble material by centrifugation at gradually increasing speed from 120×g to 27,000×g at 4 °C for 20 min. The solubilized membranes were then stored at −80 °C for future use. The protein content was determined using bicinchoninic acid (BCA) protein assay reagent (Thermo Scientific).

SDS-PAGE was performed by loading the same amount of isolated membrane proteins on a 12.5% acrylamide resolving gel. After electrophoresis, proteins were transferred to a polyvinylidene difluoride (PVDF) membrane (Millipore), blocked using 5% bovine serum albumin (BSA) for 2 h at room temperature, and then separately incubated with the primary rabbit antibodies raised against D2 and CP43 proteins overnight at 4°C. The horseradish peroxidase (HRP)-conjugated secondary antibody goat anti-rabbit IgG (H+L)-HRP conjugate (Bio-Rad) was diluted at 1:5,000 in 1.5% BSA. Bands were visualized using chemiluminescence reagents (EMD Millipore, Billerica, MA, USA) with an ImageQuant LAS-4000 imager (GE Healthcare).

### Measurement of photosynthetic activity

PSII efficiency was analyzed *in vivo* using a double-modulation fluorometer, FL-200 (Photon System Instruments, Brno, Czech Republic). 1 mL sample was dark-adapted for 2 min before measurement. The instrument contains red LEDs for both actinic (20-s) and measuring (2.5-s) flashes and was used in the time range of 100 μs to 100 s. The maximal PSII quantum yield (F_V_/F_M_) was determined with the saturation pulse method.^48^

### Cell number counting

20 μL of culture was sampled, and cells were counted with an automated cell counter (Cellometer Vision; Nexcelom). The counted images were manually curated to improve accuracy of the counts. The accompanying Cellometer software reported cell counts in cells per milliliter.

## AUTHOR INFORMATION

°D.L. and V.M.J contributed equally to this work

## Corresponding Author

*Himadri B. Pakrasi

## Author Contributions

D.L., V.M.J., and H.B.P. designed the experiments. D.L. and V.M.J. performed the experiments. D.L., V.M.J., and H.B.P. wrote the paper.

## ACKNOWLEDGMENTS

We thank members of the Pakrasi lab for congenial discussion. This work was supported by the Chemical Sciences, Geosciences, and Biosciences Division, Office of Basic Energy Sciences, Office of Science, U.S. Department of Energy (DOE) (Grant DE-FG02-99ER20350 to H.B.P.) V.M.J was supported by a training grant T32 EB014855 from the National Institute of Biomedical Imaging and Bioengineering, NIH.

