## Supplementary material for "A reversibly induced CRISPRi system targeting Photosystem II in the cyanobacterium *Synechocystis* sp. PCC 6803": SI file

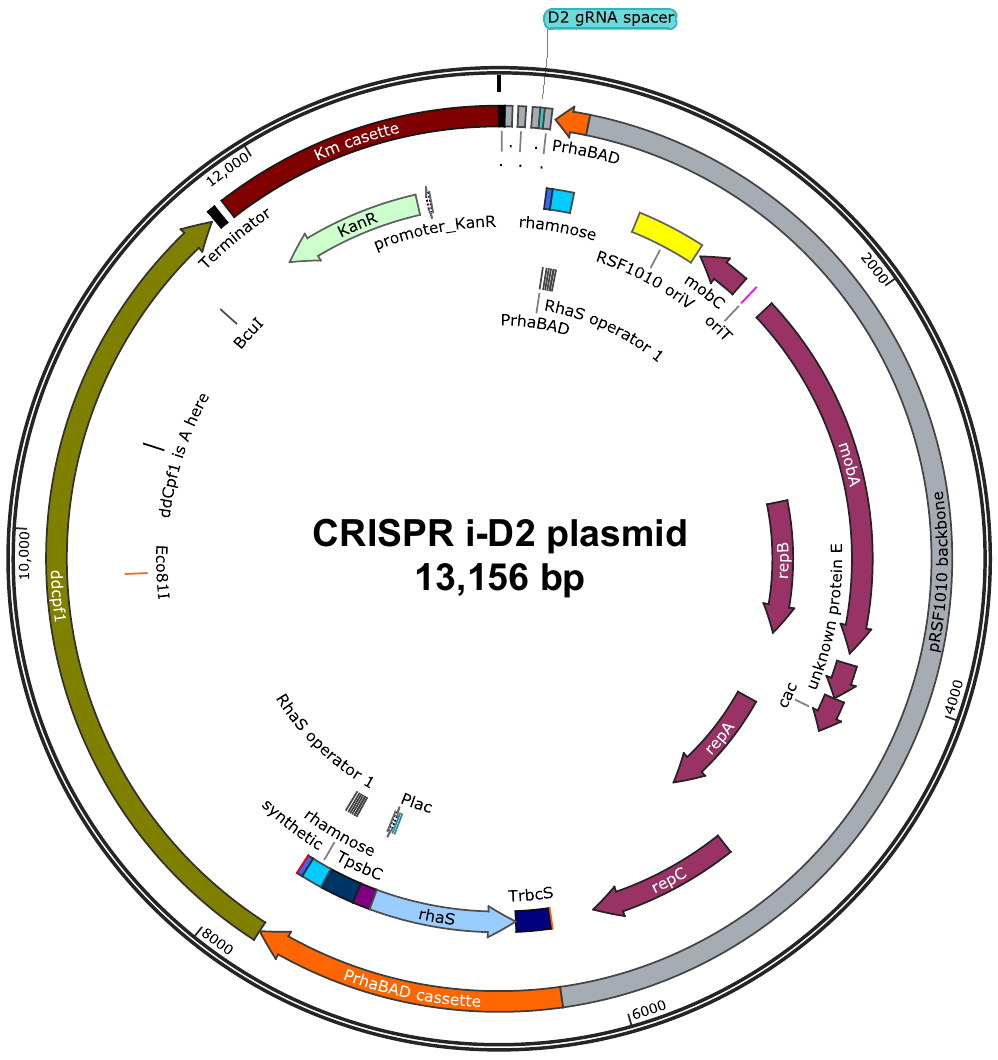

Figure S1.

Schematic map of the plasmid containing the CRISPRi system. The coding gene of regulator protein RhaS is driven by the promoter P*_lacOI_*. The sequence of the promoter P*_rhaBAD_* is identical to that in Kelly et. al.^1^ This replicating plasmid was constructed on an RSF1010 backbone using the Gibson Assembly strategy^2^ and introduced into WT of *Synechocystis* 6803 by conjugation.

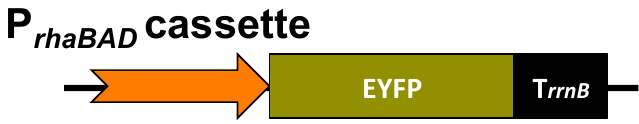

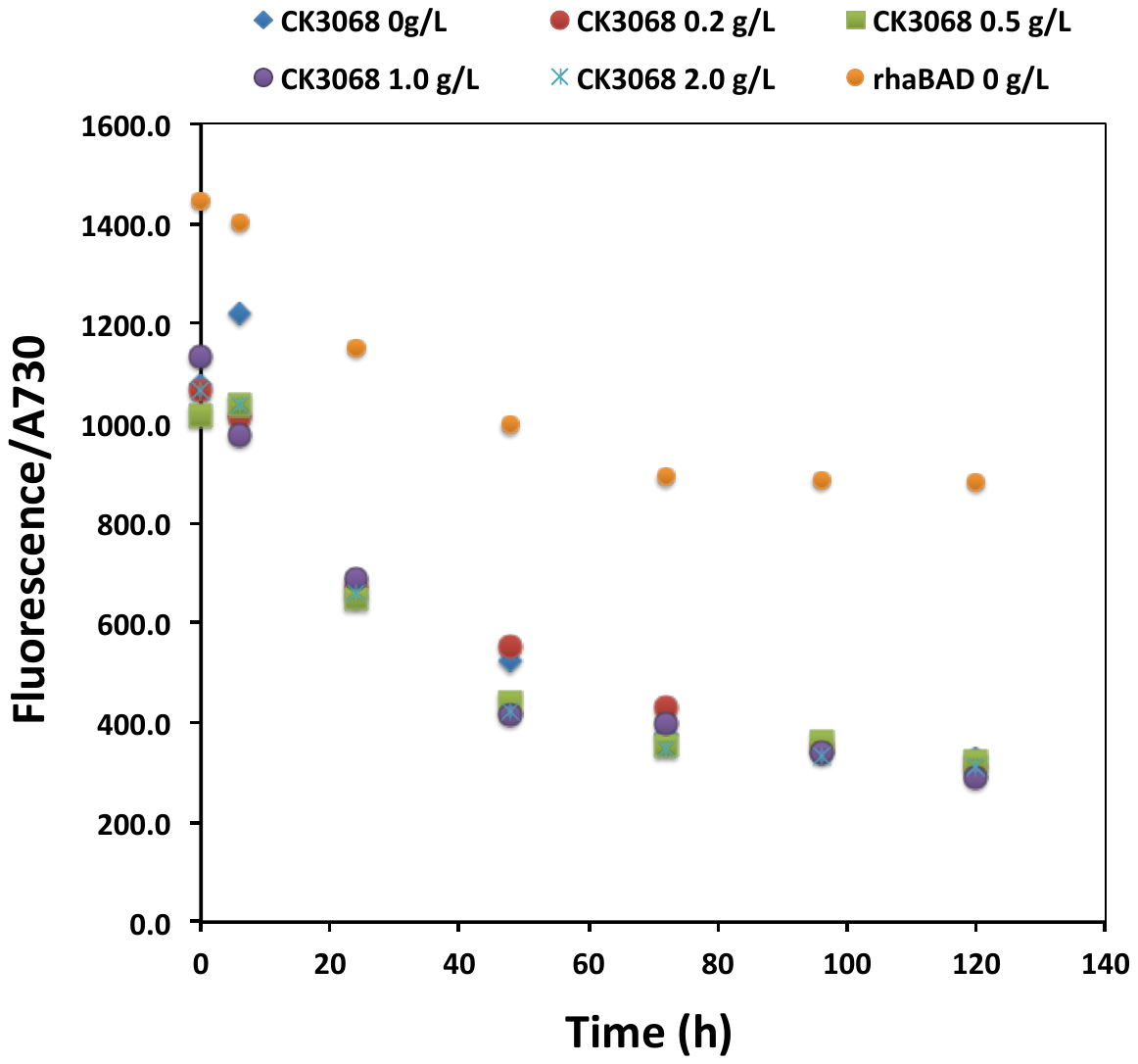

Figure S2. EYFP fluorescence levels in pRhaBAD-eyfp.

EYFP is expressed by the P*_rhaBAD_* promoter even without rhamnose added as an inducer. CK3068 is the control strain in Figure 2, which contains the plasmid with EYFP encoding gene but no promoter to drive it. Fluorescence data were normalized to optical density at 730 nm. Concentration of rhamnose added to the culture is labeled following the strain name. All strains were cultured in BG11 medium, shaking in flask under 30°C with light intensity 30 µmol photons m^−2^ s^−1^. Each dot represents the averaged data observed from at least three independent experiments.

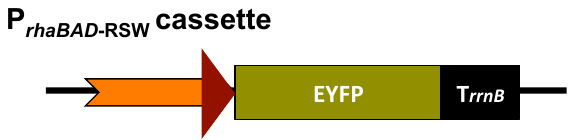

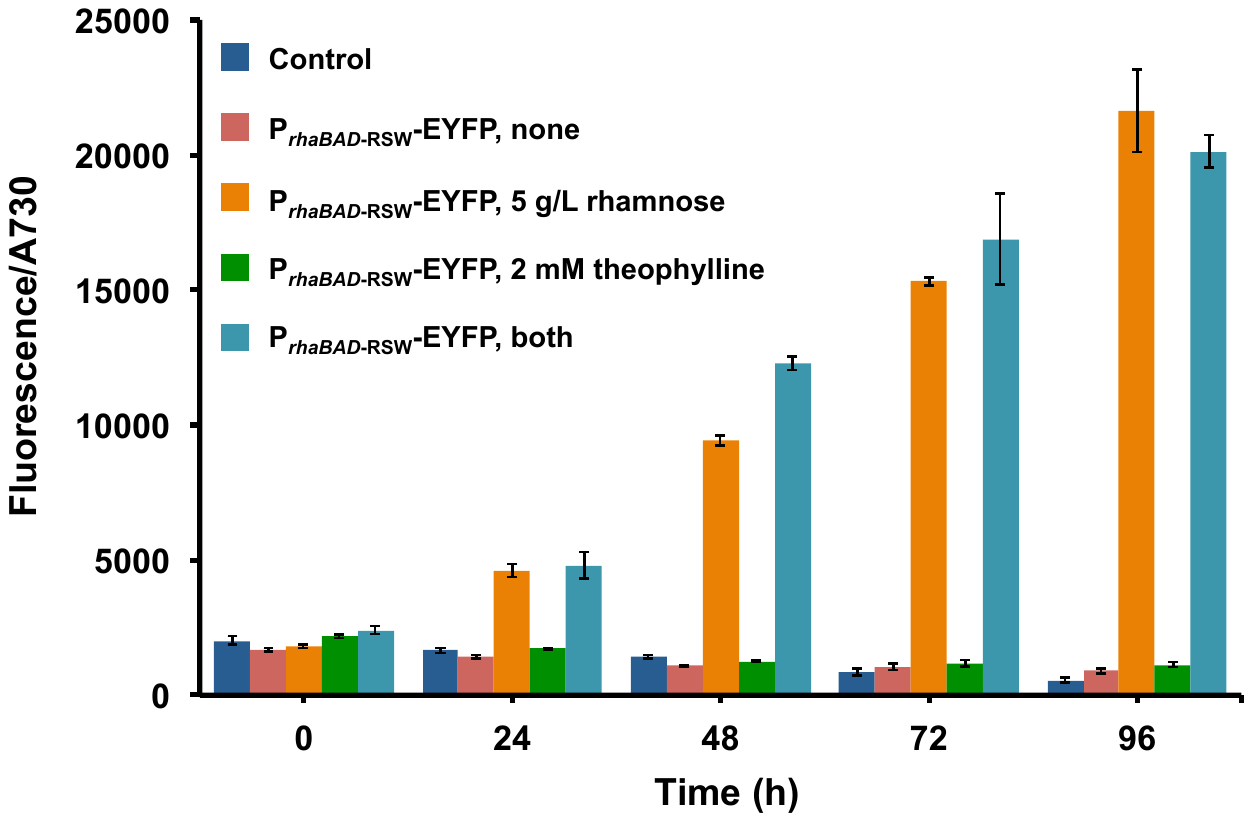

Figure S3.

Expression of EYFP driven by *P_rhaBAD_*_-RSW_, as measured with EYFP fluorescence. Control is the strain CK3068 containing the plasmid with EYFP coding gene but without any promoters to drive it. Rhamnose and theophylline were added at the concentrations indicated in the figure. “None” in the panel means no inducers were added to the culture. Fluorescence data were normalized to optical density at 730 nm. All strains were cultured in BG11 medium, shaking in flask under 30°C with light intensity 30 µmol photons m^−2^ s^−1^. Error bars represent the standard deviations observed from at least three independent experiments.

Figure S4.

Semiquantitative RT-PCR of photosystem genes. Total RNA was extracted from cell samples cultured in BG11 medium, shaking in flask under 30°C with light intensity 30 µmol photons m^−2^ s^−1^. Cells were grown with 10 g/L of glucose and 5 g/L rhamnose and 2 mM theophylline as inducers as indicated. Lane 1&5 are WT of *Synechocystis* 6803. Lane 2&6 are the control strain CK02 containing the CRISPR i_RSW_ plasmid with the gRNA targeted to the *eyfp* gene. Lane 3, 4 &7 are the CRISPR i_RSW_-D2 strain.

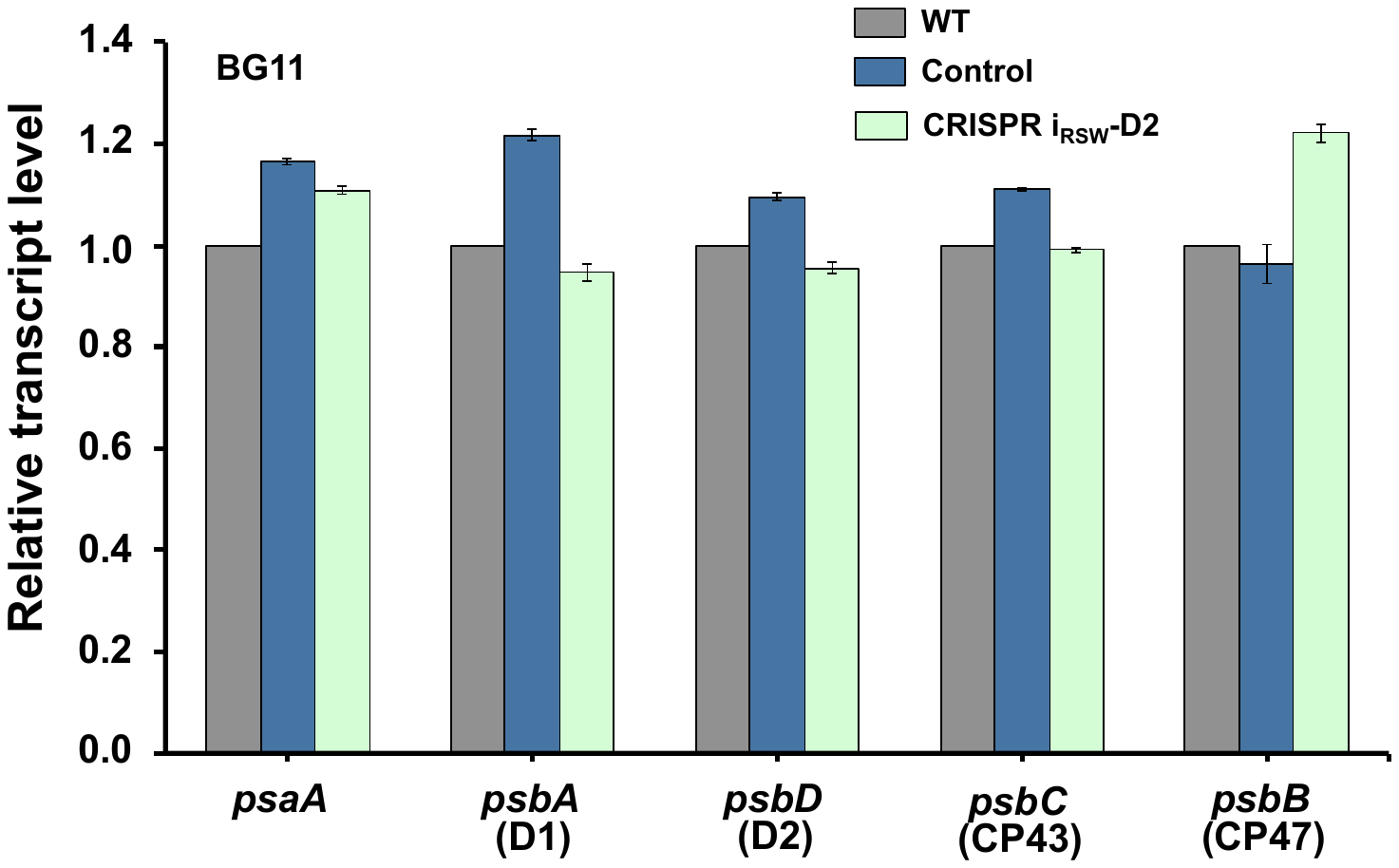

Figure S5.

Transcript levels of photosystem genes quantified by q-PCR. Cell samples were collected from culture grown in BG11 medium shaking at 30°C with 30 µmol photons m^−2^ s^−1^, and total RNA were extracted for analysis. Control is the CK02 strain containing the CRISPR i_RSW_ plasmid but with the gRNA targeting to the *eyfp* gene. Error bars represent the standard deviations observed from at least three independent experiments.

**Table S1.** Nucleotide sequences of promoters and genes in this study.

| **Promoter**  **and gene** | **Sequence** | **Note** |
| --- | --- | --- |
| P*_rhaBAD_* promoter cassette | taattgacaattgacaattccccacttagataaaaaatccggtcaggatgcggtcaacattatacaaaaactgtccaaagaacagggccgtattgtgttgctgattacctatagtcactgtgaaaaagcaacaaaaaacccgcgggagcgggctaacattgaattgaagtcagttttttagtaattgccaaaactgtaattattgcagaaagccatcccgtccctggcgaatatcacgcggtgaccagttaaactctcggcgaaaaagcgtcgaaaagtggttactgtcgctgaatccacagcgataggcgatgtcagtaacgctggcctcgctgtggcgtagcagatgtcgggctttcatcagtcgcaggcggttcaggtatcgctgaggcgtcagtcccgtttgctgcttaagctgccgatgtagcgtacgcagtgaaagagaaaattgatccgccacggcatcccaattcacctcatcggcaaaatggtcctccagccaggccagaagcaagttgagacgtgatgcgctgttttccaggttctcctgcaaactgcttttacgcagcaagagcagtaattgcataaacaagatctcgcgactggcggtcgagggtaaatcattttccccttcctgctgttccatctgtgcaaccagctgtcgcacctgctgcaatacgctgtggttaacgcgccagtgagacggatactgcccatccagctcttgtggcagcaactgattcagcccggcgagaaactgaaatcgatccggcgagcgatacagcacattggtcagacacagattatcggtatgttcatacagatgccgatcatgatcgcgtacgaaacagaccgtgccaccggtgatggtatagggctgcccattaaacacatgaatacccgtgccatgttcgacaatcacaatttcatgaaaatcatgatgatgttcaggaaaatccgcctgcgggagccggggttctatcgccacggacgcgttaccagacggaaaaaaatccacactatgtaatacggtcatagctgtttcctgtgtgaaattgttatccgctcacaattccacacaacatacgagccggaagcataaagtgtaaagcctggggtgcctaatgagtgagctaattgagacttttctgattttgcaaaggttttgctttagttaaacccaattgattagtgtcccctgcccatttggtgggggattattatttttaagataatcctattttttggagtgaggccagttacctattagacgcgcgactcgaaagtcgttcaggggagttggaacggcttccaaaaacctttccccgctggtgttccacaattcagcaaattgtgaacatcatcacgttcatctttccctggttgccaatggcccattttcctgtcagtaacgagaaggtcgcgaattcaggcgctttttagactggtcgtaatgaatcgggtaagtttataatatacaaaggaggtagaaATG | Green is the RhaS coding sequence. Blue is the P*_lacOI_* promoter. Orange is the P*_rhaBAD_* promoter. Black is the terminators. The start codon of *ddcpf1* gene is highlighted in yellow. |
| P*_rhaBAD_*_-RSW_ promoter cassette | aattgacaattgacaattccccacttagataaaaaatccggtcaggatgcggtcaacattatacaaaaactgtccaaagaacagggccgtattgtgttgctgattacctatagtcactgtgaaaaagcaacaaaaaacccgcgggagcgggctaacattgaattgaagtcagttttttagtaattgccaaaactgtaattattgcagaaagccatcccgtccctgacgaatatcacgcggtgaccagttaaactctcggcgaaaaagcgtcgaaaagtggttactgtcgctgaatccacagcgataggcgatgtcagtaacgctggcctcactgtggcgtagcagatgtcgggctttcatcagtcgcaggcggttcaggtatcgctgaggcgtcagtcccgtttgctgcttaagctgccgatgtagcgtacgcagtgaaagagaaaattgatccgccacggcatcccaattcacctcatcggcaaaatggtcctccagccaggccagaagcaagttgagacgtgatgcgctgttttccaggttctcctgcaaactgcttttacgcagcaagagcagtaattgcataaacaagatctcgcgactggcggtcgagggtaaatcattttccccttcctgctgttccatctgtgcaaccagctgtcgcacctgctgcaatacgctgtggttaacgcgccagtgagacggatactgcccatccagctcttgtggcagcaactgattcagcccggcgagaaactgaaatcgatccggcgagcgatacagcacattggtcagacacagattatcggtatgttcatacagatgccgatcatgatcgcgtacgaaacagaccgtgccaccggtgatggtatagggctgcccattaaacacatgaatacccgtgccatgttcgacaatcacaatttcatgaaaatcatgatgatgttcaggaaaatccgcctgcgggagccggggttctatcgccacggacacgttaccagacggaaaaaaatccacactatgtaatacggtcatagctgtttcctgtgtgaaattgttatccgctcacaattccacacaacatacgagccggaagcataaagtgtaaagcctggggtgcctaatgagtgagctaattgagacttttctgattttgcaaaggttttgctttagttaaacccaattgattagtgtcccctgcccatttggtgggggattattatttttaagataatcctattttttggagtgaggccagttacctattagacgcgcgactcgaaagtcgttcaggggagttggaacggcttccaaaaacctttccccgctggtgttccacaattcagcaaattgtgaacatcatcacgttcatctttccctggttgccaatggcccattttcctgtcagtaacgagaaggtcgcgaattcaggcgctttttagactggtcgtaatgaataccggtgataccagcatcgtcttgatgcccttggcagcaccctgctaaggaggtaacaacaagATG | Green is the RhaS coding sequence. Blue is the P*_lacOI_* promoter. Orange is the P*_rhaBAD_*_-RSW_ promoter. The riboswitch sequence is underlined. Black is the terminators. The start codon of *ddcpf1* gene is highlighted in yellow. |
| *psbD1* gene in *Synechocystis* 6803 | ATGACTATTGCAGTCGGACGCGCCCCAGTCGAAAGAGGATGGTTTGATGTCCTCGACGATTGGCTAAAGCGTGATCGTTTCGTATTTATTGGTTGGTCTGGTTTGCTGCTCTTCCCCTGTGCCTTCATGGCCCTGGGGGGATGGCTAACCGGCACCACCTTCGTTACTTCCTGGTACACCCACGGTCTAGCCAGTTCCTATCTAGAAGGAGCTAACTTTTTGACCGTGGCGGTCTCTTCCCCCGCCGATGCCTTCGGCCATTCCCTCCTGTTCCTCTGGGGACCCGAAGCCCAAGGTAACCTGACCCGCTGGTTCCAAATCGGTGGTTTGTGGCCCTTCGTTGCCCTCCACGGTGCCTTCGGTCTGATTGGCTTTATGCTGCGTCAGTTCGAAATTTCCCGTCTGGTTGGCATTCGTCCCTACAACGCCATCGCCTTCTCTGGTCCCATTGCGGTGTTTGTCAGTGTCTTTTTGATGTACCCCTTGGGACAATCCAGTTGGTTCTTTGCCCCCAGCTTTGGGGTAGCGGGAATCTTCCGGTTCATTTTGTTCCTGCAAGGGTTCCACAACTGGACTTTGAACCCCTTCCACATGATGGGAGTGGCAGGGATTTTGGGCGGCGCCCTCCTCTGTGCTATCCACGGTGCCACGGTGGAAAACACCCTGTTTGAAGATGGGGAAGATTCCAATACTTTCCGGGCATTTGAACCCACCCAAGCAGAAGAAACCTATTCCATGGTGACCGCTAACCGTTTCTGGTCTCAGATTTTCGGTATTGCTTTCTCCAACAAGCGGTGGTTGCACTTCTTCATGTTGTTCGTGCCGGTAACTGGTCTGTGGATGAGTTCCGTCGGTATCGTCGGTTTAGCCTTGAACCTACGGGCCTATGACTTTGTCTCCCAGGAGCTACGGGCGGCGGAAGACCCGGAATTTGAAACTTTCTATACGAAAAACATTTTGTTGAACGAAGGGATGCGCGCCTGGATGGCTCCCCAAGATCAACCCCATGAAAACTTTATCTTCCCTGAGGAGGTTCTCCCCCGTGGTAACGCTCTCTAA | Red is the sequence for gRNA recognition |
| *psbD2* gene in *Synechocystis* 6803 | ATGACCATTGCAGTCGGACGCGCCCCAGTCGAAAGAGGATGGTTTGATGTCCTCGACGATTGGCTAAAGCGTGATCGTTTCGTATTTATCGGTTGGTCTGGTTTGCTACTCTTCCCCTGCGCCTTCATGGCCCTGGGGGGATGGTTAACCGGCACCACCTTCGTTACTTCCTGGTACACCCACGGTCTAGCCAGTTCCTACCTGGAAGGGGCTAACTTTTTGACCGTGGCGGTCTCTTCCCCCGCCGATGCCTTCGGCCATTCCCTCCTGTTCCTGTGGGGACCGGAAGCTCAAGGTAACCTGACCCGCTGGTTCCAAATTGGTGGTTTGTGGCCCTTCGTTGCCCTCCACGGTGCCTTTGGATTGATTGGCTTCATGCTGCGTCAGTTCGAAATTTCCCGTCTGGTAGGCATTCGTCCCTACAACGCCATCGCTTTCTCTGGTCCCATTGCGGTATTTGTCAGCGTCTTTCTGATGTACCCCTTGGGTCAATCGAGTTGGTTCTTTGCTCCCAGCTTTGGGGTAGCGGGAATCTTCCGGTTTATTTTGTTCCTACAAGGTTTCCACAACTGGACCCTGAACCCCTTCCACATGATGGGAGTAGCCGGTATTCTCGGTGGTGCCCTACTGTGTGCCATCCACGGTGCCACGGTGGAAAACACCCTGTTTGAAGACGGTGAAGATTCCAACACCTTCCGGGCGTTTGAACCTACCCAAGCGGAAGAAACCTACTCCATGGTGACTGCCAACCGTTTCTGGTCTCAGATTTTCGGTATTGCTTTCTCCAACAAACGGTGGCTGCACTTCTTCATGTTGTTCGTTCCCGTAACTGGTTTGTGGATGAGTTCTGTGGGTATCGTCGGTTTGGCGTTGAACCTACGGGCTTATGACTTCGTTTCCCAGGAACTGCGGGCTGCTGAAGATCCGGAATTTGAAACGTTTTATACGAAAAACATTTTGTTGAACGAAGGGATGCGCGCCTGGATGGCTCCCCAAGATCAACCCCATGAAAACTTTATCTTCCCTGAGGAAGTACTGCCCCGGGGTAATGCTCTCTAA | Red is the sequence for gRNA recognition |

**Table S2.** Strains and plasmids used in this study.

| **Strains and plasmids** | **Description** | **Source** |
| --- | --- | --- |
| **Strain** |  |  |
| *E.coli* strain XL1-Blue | Used for DNA cloning | Pakrasi lab |
| *Synechocystis* 6803 | Wild type (WT) strain | Pakrasi lab |
| CK3068 | Control strain containing the eyfp gene on pRSF1010 but without any promoters to drive it | 3 |
| CK01 | Control strain containing the CRISPRi system but gRNA targeted to *eyfp* | This study |
| CK02 | Control strain containing the CRISPR i_RSW_ system but targeted to *eyfp* by the gRNA | This study |
| P*_rhaBAD_*-EYFP | Strain containing the eyfp gene on pRSF1010 driven by the P*_rhaBAD_* promoter | This study |
| P*_rhaBAD_*_-RSW_-EYFP | Strain containing the eyfp gene on pRSF1010 driven by the P*_rhaBAD_*_–RSW_ promoter | This study |
| CRISPR i-D2 | Strain containing the CRISPRi system with gRNA targeted to *psbD* genes of *Synechocystis* 6803 | This study |
| CRISPR i_RSW_-D2 | Strain containing the CRISPR i_RSW_ system with gRNA targeted to *psbD* genes of *Synechocystis* 6803 | This study |
| **Plasmid** |  |  |
| pRL443 | Plasmid used for conjugation | 4 |
| PRL623 | Plasmid used for conjugation | 5 |
| pRSF1010 | Broad-host-range shuttle vector | 6 |
| pCK3068 | Plasmid containing the eyfp gene on pRSF1010 but without any promoters to drive it | 3 |
| pCRISPR i-eyfp | Plasmid containing CRISPRi system on pRSF1010 with gRNA targeted to *eyfp* gene | This study |
| pCRISPR i_RSW_-eyfp | Plasmid containing CRISPR i_RSW_ system on pRSF1010 with gRNA targeted to *eyfp* gene | This study |
| pRhaBAD-eyfp | Plasmid containing the *eyfp* gene on pRSF1010 driven by P*_rhaBAD_* | This study |
| pRhaBAD-RSW-eyfp | Plasmid containing the *eyfp* gene on pRSF1010 driven by P*_rhaBAD_*_-RSW_ | This study |
| pCRISPR i-D2 | Plasmid containing the CRISPRi system on pRSF1010 with gRNA targeted to *psbD* gene of *Synechocystis* 6803 | This study |
| pCRISPR i_RSW_-D2 | Plasmid containing the CRISPR i_RSW_ system on pRSF1010 with gRNA targeted to *psbD* gene of *Synechocystis* 6803 | This study |

**Table S3** List of primers used in this study.

| **Primer name** | **Sequence 5' -> 3'** | **Purpose of primers** |
| --- | --- | --- |
| RHA_TrbcS_F1 | cgctttcctggctttgcttcccactaattgacaattgacaattccccac | For plasmid pRhaBAD-eyfp |
| RHA_TrbcS_R1 | acgggatggctttctgcaataattacagttttggcaattactaaaaaac |  |
| RHA_rhaS_F2 | gccaaaactgtaattattgcagaaagccatcccg |  |
| RHA_rhas_R2 | caggaaacagctatgaccgtattacatagtgtgg |  |
| RHA_PlacOI_F3 | gtaatacggtcatagctgtttcctgtgtgaaattg |  |
| RHA_PlacOI_R3 | aagtctcaattagctcactcattaggcacccc |  |
| RHA_TpsbC_F4 | gagtgagctaattgagacttttctgattttgc |  |
| RHA_TpsbC_R4 | tgaattgtggaacaccagcggggaaaggtttttg |  |
| RHA_Prha_F5 | cgctggtgttccacaattcagcaaattgtgaac |  |
| RHA-Prha_R5 | attataaacttacccgattcattacgaccagtctaaaaagc |  |
| RHA_Prha_R6 | ttctacctcctttgtatattataaacttacccgattcattac |  |
| RHA_Prha_R7 | cagctcctcgcccttgctcaccatttctacctcctttgtatattataaac |  |
| EYFP_F | atggtgagcaagggcgaggag |  |
| TrrnB_R | cattacgctgacttgacgggacacggttttaaagaaaaagggcagg |  |
| PrhaBAD-RSW_KspAI_F1 | ctgtcgcacctgctgcaatac | For plasmid pRhaBAD-RSW-eyfp |
| PrhaBAD-RSW_R1 | agggcatcaagacgatgctggtatcaccggtattcattacgaccagtctaaaaagcgc |  |
| PrhaBAD_RSW_R2 | tgttgttacctccttagcagggtgctgccaagggcatcaagacgatgctgg |  |
| PrhaBAD-RSW_F3 | tggcagcaccctgctaaggaggtaacaacaagatggtgagcaagggcgaggag |  |
| PrhaBAD-RSW_pstI_R4 | aaagggcagggtggtgacac |  |
| Rha_lguI_F1 | ctggctttgcttccagatgtatgctcttcttaattgacaattgacaattccccac |  |
| PrhaBAD_F | ccgtttttgcctaaatcagcttctacctcctttgtatattataaac | For plasmid pCRISPR i-eyfp |
| PrhaBAD_R | gagtcttgccacgccgagcacctggccacaattcagcaaattgtgaac |  |
| Template2_R | gctgatttaggcaaaaacgg |  |
| Template1_F | cgtggctttgcgcattaaagc |  |
| Template_F3 | tacagctgaaagcgaaaaaaaatgaccttcataaatcgc |  |
| cpf1_BcuI_F | agagcgacaaaaagttttttgc |  |
| cpf1_BcuI_R | aaggtcatttttttgtctagc |  |
| Km_F2 | gctagacaaaaaaatgacctttcgactctagagaggatctgtaac |  |
| Km_R2 | gaaggtcattttttttcgctttcagctgtaatccgg |  |
| leader-rha_R | ggaggtagaagctgatttaggcaaaaacgg |  |
| Rha_RSW-dcpf1_R1 | tttattaacaaattcttgataaattgacatcttgttgttacctccttagcagg | For plasmid pCRISPR i_RSW_-eyfp |
| dcpf1_start_F2 | atgtcaatttatcaagaatttgttaataaatatag |  |
| dCpf1_lguI_R2 | tgtagtcaatatcaaaggttagctc |  |
| EcoO109I_R | gatgagccgggctgaatgatc | For plasmid pCRISPR i-D2 |
| OliI_F | agacctcagcgctattctgac |  |
| Rhacasette_F | atgtctagctttaatgcggtagttagatcttaattgacaattgacaattccccac |  |
| RHA_PlacOI_F3 | gtaatacggtcatagctgtttcctgtgtgaaattg |  |
| Rhacasette_R | tgataaattgacatttctacctcctttgtatattataaacttac |  |
| qPCR_6803RnpB_F0 | agtatcgagaggtactggctc | For q-PCR |
| qPCR_6803RnpB_R0 | acccttggggagttatctatc |  |
| qPCR_6803D1_F1 | ctctaccaacaaccggatttatg |  |
| qPCR_6803D1_R1 | gcgatgaaggcaatgatgaag |  |
| qPCR_6803D2_F2 | ttacttcctggtacacccacg |  |
| qPCR_6803D2_R2 | gaaccagcgggtcaggttac |  |
| qPCR_6803CP43_F3 | ccgtggtaacgctctctaatac |  |
| qPCR_6803CP43_R3 | gattaatcagccgggcatttc |  |
| qPCR_6803CP47_F4 | ccttggtatcgcgttcatacag |  |
| qPCR_6803CP47_R4 | gccaactcatagagagccatag |  |
| qPCR_6803psaA_F5 | gataccgctcaccaccatttg |  |
| qPCR_6803psaA_R5 | cgaggatctctttcatgctatgg |  |
